# Pandemic Response Box Screening Identified CRM1/XPO1 as an Anti-Mammarenavirus Druggable Target

**DOI:** 10.1101/2025.11.24.690217

**Authors:** Chukwudi A. Ofodile, Beatrice Cubitt, Ngozi Onyemelukwe, Chetachi B. Okwuanaso, Haydar Witwit, Juan C. de la Torre

## Abstract

Mammarenaviruses (MaAv) cause persistent infection in their natural rodent hosts across the world and via zoonotic events can cause severe disease in humans. Thus, the MaAv Lassa virus (LASV) in Western Africa and Junin virus (JUNV) in the Argentinean Pampas cause hemorrhagic fever diseases with significant case fatality rates in their endemic regions. In addition, the globally distributed MaAv lymphocytic choriomeningitis virus (LCMV) is an underrecognized human pathogen of clinical significance capable of causing devastating infections in neonates and immunocompromised individuals. Despite their impact on human health, there are currently no FDA-approved vaccines or specific antiviral treatments for MaAv infections. Existing anti-MaAv therapies are limited to the off-label use of ribavirin whose efficacy remains controversial, hence, the significance of developing novel therapeutics to combat human pathogenic MaAv. We employed a high-throughput cell-based infection assay to screen the Pandemic Response Box, a collection of 400 diverse compounds with established antimicrobial activity, for MaAv inhibitors. We identified Ro-24-7429, an antagonist of the HIV-1 Tat protein and RUNX Family Transcription Factor 1 inhibitor; WO 2006118607 A2, a dihydroorotate dehydrogenase inhibitor; and verdinexor, a novel selective inhibitor of nuclear export (SINE) targeting the CRM1/XPO1, as potent anti-MaAv compounds. Consistent with their distinct validated targets, verdinexor and WO 2006118607 A2 exhibited a very strong synergistic antiviral activity when used in combination therapy. Our findings pave the way for the development of verdinexor as a potent host-directed antiviral against MaAv, which could be integrated into the development of combination therapy with direct-or host-acting antivirals to combat human pathogenic MaAv.

## 1. Introduction

Mammarenaviruses (MaAv) cause persistent infections in their natural rodent reservoirs worldwide [1]. Human infections occur through mucosal exposure to aerosols or by direct contact of abraded skin with infectious materials[1]. Several MaAv cause hemorrhagic fever (HF) in humans and represent important public health concerns in their endemic regions [1]. Lassa virus (LASV), endemic to Western Africa, is the etiologic agent of Lassa fever (LF), a HF disease associated with high morbidity and a case fatality rate as high as 69% among hospitalized confirmed LF cases [2–5]. LASV is estimated to infect >500,000 people annually [6], but limited diagnostic capabilities complicates an accurate assessment of LASV prevalence [7,8]. Recent studies indicate that LASV endemic regions are expanding [9–11], and increased traveling has led to the importation of LF cases into non-endemic metropolitan areas around the globe [12,13]. Notably, LASV ranks very high among zoonotic viruses with potential for spillover and spread in humans [14] and models based on projections of climate, land use and population changes indicate that regions in Central and East Africa will become suitable for LASV over the next decades resulting in a dramatic increase in the population at risk of LASV exposure [15]. Because of its impact in human health, pandemic potential and lack of effective countermeasures, LF has been included within the World Health Organization (WHO) list of priority diseases for which there is an urgent need to develop therapeutic and vaccines [16].

There are no US FDA-approved MaAv vaccines and current anti-MaAv therapy is limited to an off-label use of ribavirin whose efficacy remains controversial [17,18]. The broad-spectrum polymerase inhibitor favipiravir [19,20] and the cell entry inhibitors LHF-535 [21] and ARN-75039 [22,23] have shown promising results in preclinical models of LF, and LHF-535 is in phase I clinical trials. However, favipiravir limited potency against LASV and short human half-life [24,25] are likely to complicate its efficacy. Likewise, the high degree of LASV genetic diversity [26,27] compromises the use of cell entry inhibitors as single mutations within the SSP or the TMD can confer cross-resistance to these cell entry inhibitors [28,29]. The development of broad-spectrum oral antivirals to counteract human pathogenic MaAv represents a significant unmet biomedical need.

MaAv are enveloped viruses with a bisegmented, negative strand RNA genome [1,30,31]. Each genome segment uses an ambisense coding strategy to express two viral proteins. The large (L) segment (ca. 7.3 kb) encodes the RING-finger matrix protein (Z), and the RNA-dependent RNA polymerase (RdRP) L protein. The small (S) segment (ca 3.4 kb) encodes the glycoprotein precursor (GPC) and nucleoprotein (NP). NP encapsulates the viral genome RNA to create the viral nucleocapsid (NC) that together with the L protein form the viral ribonucleoprotein complex (vRNP) that directs replication and transcription of the viral genome [1,32]. GPC is co-translationally processed by the signal peptidase to form a stable signal peptide (SSP) and a GPC precursor, which is post-translationally processed by the S1P protease into GP1 and GP2. The mature tripartite GP complex, consisting of SSP, GP1, and GP2 decorates the surface of the virion and mediates virus cell entry via receptor-mediated endocytosis [32]. GP1 binds to cellular receptors, whereas GP2 mediates a pH-dependent fusion event between viral and cell membranes in the acidic environment of the late endosome [33]. Upon fusion, the vRNP is released into the cytosol, where it initiates viral transcription and replication [1]. MaAv assembly and budding is driven by the matrix protein Z [1].

Drug repurposing strategies can substantially decrease the time and resources necessary to advance a candidate antiviral drug into the clinic [34]. Furthermore, drug repurposing can yield valuable insights into viral biology by identifying novel pathways and host cell factors involved in various stages of the viral replication cycle, thereby revealing new antiviral targets [35–38].

Here, we document the results of the screening of the Pandemic Response Box (PRB) [39] for inhibitors of MaAv infection. The PRB was assembled by the Medicines for Malaria Venture, Geneva (MMV) and Drugs for Neglected Diseases initiative, (DNDi) and contains a collection of 400 compounds comprising 201 antibacterial, 153 antiviral, and 46 antifungal compounds, with several of these compounds demonstrating activity across multiple disease domains [39]. We identified three compounds listed in the PRB as Ro-24-7429 (RO), WO 2006118607 A2 (WO), and Verdinexor (VE) as potent inhibitors of the activity of LCMV and LASV vRNP in cell-based minigenome assays, and multiplication of LCMV in cultured cells.

RO is an antagonist of the HIV-1transcriptional transactivator (Tat) protein that inhibits HIV-1 replication via interference with Tat-dependent initiation and elongation of transcription, resulting in reduced levels of viral RNA synthesis and protein production [40,41] but with no detectable antiviral effect [42]. RO inhibits RUNX Family Transcription Factor 1, which has been associated with its antifibrotic and anti-inflammatory properties[43]. WO has been shown to have antiviral activity against HCV via inhibition of dihydroorotate dehydrogenase (DHODH), a key enzyme in pyrimidine biosynthesis [44,45]. VE is a novel oral Selective Inhibitor of Nuclear Export (SINE) compound that targets exportin 1 (XPO1) [46]. VE has been shown to inhibit multiplication of IAV [47] and RSV [48] in cultured cells, and has shown efficacy in animal models of IAV infection [47,49]. RO, WO and VE target different distinct host, rather than viral, factors. Consistent with their distinct validated targets, VE and WO exhibited a very strong synergistic activity when used in combination therapy. Host-directed antivirals (HDAs) represent a promising yet underexplored approach in contemporary antiviral research [50]. Unlike conventional direct-acting antivirals (DAAs) that target viral proteins and functions, HDAs disrupt host factors and cellular processes that viruses hijack for their replication and pathogenesis. Since members of a virus family share host dependencies, HDAs have the potential to act as broad-spectrum antivirals. In addition, HDAs pose a higher genetic barrier to the emergence of drug-resistant viral variants, which often compromise antiviral therapy [51]. Likewise, combination therapy of antivirals with synergistic effects can pose a high genetic barrier to the emergence of drug-resistant viral variants that often jeopardize monotherapy approaches and facilitate the use of reduced drug doses within therapeutic range to alleviate side effects associated with high drug doses used in monotherapy.

## 2. Materials and Methods

### 2.1 Compound source

Medicines for Malaria Venture, Geneva (MMV) and Drugs for Neglected Diseases initiative, Geneva, Switzerland (DNDi) assembled the Pandemic Response Box (PRB), a collection of 400 compounds [39].

### 2.2 Cells and viruses

HEK 293T (ATCC CRL-3216), Vero E6 (ATCC CRL-1586), and A549 (ATCC CCL-185) cells were cultured in Dulbecco’s modified Eagle’s medium (DMEM; Thermo Fisher Scientific, Vacaville, CA, USA). The medium was supplemented with 10% heat-inactivated fetal bovine serum (FBS), 2 mM L-glutamine, and antibiotics (100 µg/mL streptomycin and 100 U/mL penicillin). Recombinant viruses rLCMV/GFP-P2A-NP[52], rARMΔGPC/ZsG-P2A-NP [53] have been described. Using the same strategy described for the generation of rLCMV/GFP-P2A-NP, we generated a rJUNV/GFP-P2A-NP based on the live-attenuated vaccine strain Candid 1 of JUNV.

### 2.3 Generation of rJUNV/GFP-P2A-NP

For the rescue of rJUNV/GFP-P2A-NP, based on the live-attenuated vaccine strain Candid 1 of JUNV, we used the same procedures and strategy described for the generation of rLCMV/GFP-P2A-NP[52]. Briefly, BHK-21 cells were seeded into 6-well plates at 7.0 × 10^5^/well 1-day prior transfection to achieve an 80-90% confluent. Cells were transfected with 1.2 µg of pC-NP and 1.5 µg of pC-L (both NP and L corresponding to the Romero strain of JUNV), 1.2 µg of mouse pol-I S JUN GFP-P2A-NP and 2.1 µg of the mouse pol-I L JUNV (total 6 µg/well) using Lipofectamine 3000 (LF3000) (Invitrogen) (1.25 µl LF3000/ µg of plasmid DNA) and P3000™ reagent (1.75 µl P3000/ µg of plasmid DNA) for more efficient transfection. At 72 h post-transfection, cell culture supernatants (CCS) were collected, and virus titration was performed.

### 2.4 Virus titration

Virus titers were determined by a focus-forming assay (FFA) [54] using Vero E6 cells. Briefly, cells were seeded in a 96-well (2.0 × 10^4^ cells/well) and 24 h later infected with 10-fold serial dilutions of the virus. At 20 h post-infection (hpi), the cells were fixed with 4% paraformaldehyde (PFA) in PBS, and infected cells (detected by epifluorescence through their expression of GFP).

### 2.5 Cell viability assay

Cell viability was evaluated by measuring formazan produced from MTS [3-(4,5-di-methylthiazol-2-yl)-5-(3-carboxymethoxyphenyl)-2-(4-sulfophenyl)-2H-tetrazolium] through the activity of NADPH or NADH produced by living cells. Cells were seeded in a 96-well optical plate (3.0 × 10^4^ cells/well) and 20 hours later, compounds (at designated concentrations, four replicates) or vehicle control (0.3% DMSO) were added, bringing the final volume to 100 µL. After 72 h treatment, the CellTiter 96 AQueous One Solution reagent (Promega, CAT#: G3580, Madison, WI, USA) was added to each well, and the plate incubated for 15 minutes at 37 °C with 5% CO2. Absorbance was measured at 490 nm using Biotek Cytation 5 plate reader (Agilent Technologies, Santa Clara, CA, USA). The results were normalized to the vehicle control which was assigned a value of 100%. One Solution and media were then aspirated, and the cells fixed with 4% paraformaldehyde and stained with DAPI.

### 2.6 Determination of compounds EC50 and CC50 values

Cells were seeded onto a 96-well optical plate (3.0 × 10^4^ cells/well), and 20 hours later cells were infected with rLCMV/GFP-P2A-NP at MOI of 0.03 or rJUNV/GFP-P2A-NP at MOI of 0.03. Infected cells were treated with three-fold serial dilutions of indicated compound, starting at a concentration of 30 µM and using four replicates for each dilution. After 72 h treatment, EC50 values were determined by measuring GFP expression levels. After measuring GFP, cells were subjected to DAPI staining. GFP and DAPI were quantified by fluorescence using a Biotek Cytation 5 plate reader (Agilent Technologies, Santa Clara, CA, USA). Mean values were normalized to an infected and vehicle (DMSO)-treated control, also set to 100%. The EC50 and CC50 values were calculated using GraphPad Prism v10 (GraphPad Software, San Diego, CA, USA).

### 2.7 Virus multi-step growth kinetics

A549 cells were seeded (2.0 × 10^5^ cells/well, a 12-well plate) and the following day infected with rLCMV/GFP-P2A-NP at an MOI of 0.03. After a 90-minute adsorption period, the inoculum was removed, and the cells were treated with VC (0.3% DMSO), specified compounds (1 µM and 5 µM), or ribavirin (RBV)(100 µM) as a positive control. CCS were collected at 24, 48, and 72 hpi. At 72 hpi, cells were stained with Hoechst dye solution. Live cell images were taken using a Keyence BZ-X710 all-in-one fluorescence microscope series. Cells were washed, and RNA was extracted using TRI Reagent (TR 118, Molecular Research Centre, Cincinnati, OH, USA). Virus titers in CCS were measured by FFA using Vero E6 cells (4 replicates) and results expressed as mean ± SD. Data were plotted using GraphPad Prism (GraphPad Software, San Diego, CA, USA).

### 2.8 RT-qPCR

RT-qPCR was conducted as described [55]. RNA isolation was done using TRI Reagent following the manufacturer’s guidelines. RNA was dissolved in RNA storage solution (Life Technologies, Carlsbad, CA, USA. AM 7000) and its concentration was measured with a NanoDrop™ 2000 spectrophotometer (ND-2000 Thermo Fisher Scientific™). RNA (1 µg) was converted into cDNA using the SuperScript IV first-strand synthesis system (18091050, Life Technologies). For amplification of LCMV NP, and the housekeeping gene GAPDH, Powerup SYBR (A25742, Life Technologies) was employed, utilizing the following primers: NP forward (F): 5′ CAGAAATGTTGATGCTGGACTGC-3′ and NP re-verse (R): 5′-CAGACCTTGGCTTGCTTTACACAG-3′; GAPDH F: 5′-CATGA-GAAGTATGACAACAGCC-3′ and GAPDH R: 5′-TGAGTCCTTCCACGATACC-3′; ISG15 F: 5′-CAGGACGACCTGTTCTGGC-3′ and ISG15 R: 5′-GATTCATGAACAC-GGTGCTCAGG-3′; and MX1 F: 5′-GCAGCTTCAGAAGGCCATGC-3′ and MX1 R: 5′-CCTTCAGGAACTTCCGCTTGTC-3′.

### 2.9 Time of addition assay

A549 cells were seeded (3.0 × 10^4^ cells/well) in a 96-well black-walled optical plate (353219, Falcon) and after 16 h culture treated with VC (0.3% DMSO), indicated compounds at concentrations of 1 µM and 5 µM, and F3406 at 5 µM, with four replicates for each treatment condition. Treatments were administered either 2 hours prior (-2h) or 2 hours after (+2h) infection with the single-cycle infectious rARMΔGPC/ZsG-P2A-NP (MOI of 1.0), thereby obviating the need for NH4Cl treatment to avoid confounding effects from multiple rounds of infection. At 24 hpi, cells were fixed with 4% PFA, and ZsGreen expression levels were quantified using a Cytation 5 imaging reader.

### 2.10 Budding Assay

Budding assays were done as described [56]. Briefly, HEK293T cells (1.75 × 10^5^ cells/well in 12-well plate) were transfected with 0.5 µg of pC.LASV-ZGluc or pC.LASV-Z-G2A-GLuc (mutant control) or pCAGGS-Empty (pC-E) using Lipofectamine 3000 and P3000™. After 5 h transfection, cells were washed three times and treated with VE, RO, WO (1 µM and 5 µM), RBV (100 µM), or vehicle control (VC). After 48 h of treatment, CCS containing virus-like particles (VLPs) were collected and cell lysates prepared. CCS samples were clarified from cell debris by centrifugation (13,000 rpm/4 °C/10 min) and aliquots (20 µL each) from CCS samples were added to 96-well black plates (VWR, West Chester, PA, USA) and 50 µL of Steady-Glo luciferase reagent (Promega) added to each well. Cell lysates were prepared using 250 µL of lysis buffer (1% NP-40, 50 mM Tris-HCl (pH 8.0), 62.5 mM EDTA, 0.4% sodium deoxycholate). Lysates were clarified from cell debris by centrifugation (13,000 rpm/4 °C/10 min). GLuc activity in Z-containing VLP and whole-cell lysates (WCLs) was determined using the Steady-Glo luciferase assay system (Promega, Madison, WI, USA) according to the manufacturer’s protocol using a Berthold Centro LB 960 luminometer (Berthold Technologies, Oak Ridge, TN, USA). The GLuc activity in CCS and WCL served as a surrogate for Z protein levels. Z budding efficiency (in %) was determined by the ratio of VLP-associated GLuc levels (ZVLP) and total GLuc levels (ZVLP + ZWCL) times 100.

### 2.11 LCMV and LASV cell-based minigenome (MG) assays

The LCMV and LASV MG systems have been described [57,58]. HEK293T cells were seeded (0.75 × 10^6^ cells/well) onto a poly-L-lysine-treated 6-well plate 20 hours prior to transfection with pCAGGS-T7 Cyt (0.625 µg), and the components (pT7MG-ZsGreen, pCAGGS-NP, and pCAGGS-L. Cells were transfected using Lipofectamine 3000™ (2.5 µL/2.4 µg DNA) and P3000™ (4 µL/2.4 µg DNA). After 2.5 h, the transfection medium was removed, and the cells were washed with DPBS. Cells were detached using 0.5 mL of Accutase, collected with 1 mL of fresh medium, and centrifuged at 1300 rpm at 6°C for 5 minutes. The transfected cells were resuspended in an appropriate volume of fresh medium to achieve a cell density of 3.0 × 10^4^ cells per well in a 96-well plate. Subsequently, 10 µL (10x) of three-fold serial dilutions of the indicated compounds, starting at 20 µM (VE and RO), and 1 µM (WO) were prepared in four replicates and plated onto a 96-well plate. Then, 90 µL of the cell suspension (3.0 × 10^4^ cells/well) was added, bringing the final volume to 100 µL, and incubated for 72 hours at 37°C with 5% CO2. The cells were then fixed with 4% paraformaldehyde and stained with DAPI. EC50 values were determined by measuring GFP expression levels, while CC50 was evaluated by measuring DAPI. Both GFP and DAPI were quantified by fluorescence using a Biotek Cytation 5 plate reader (Agilent Technologies, Santa Clara, CA, USA). The mean values were normalized to the vehicle (DMSO)-treated control, which was set to 100%. The LASV L and NP sequences used in the MG assay were derived from the Josiah strain.

### 2.12 Epifluorescence

Images were collected using the Keyence BZ-X710 microscope. Images were transferred to a laptop for data processing purposes. Microsoft PowerPoint Version 16.103 (25110922) was used to assemble and arrange the images, with each one being imported separately and arranged in a cohesive manner within its respective composite. The canvas size was adjusted to ensure a harmonious layout.

### 2.13 siRNA-mediated XPO1 Knockdown (KD)

ON-TARGETplus Human XPO1 (7514) siRNA – SMARTpool (Dharmacon) with four target sequences: “AAGAAUGGCUCAAGAAGUA”, “GGACAA-GAGUCGACACAAU”, “UAGAUAAUGUGGUGAAUUG” and “CGAAAUGUCU-CUC” was used to KD XPO1 in A549 cells using non-targeting siRNA as control. Briefly, A549 cells were seeded overnight in 12 well plate at 2.0 × 10^5^/well and transfected with siRNA targeting XPO1 and non-targeting siRNA control using DharmaFECT-1 reagent (Dharmacon). 10μl of 5μM siRNA was diluted with 190μl of OptiMem to give a concentration of 250 nM of siRNA. 2 μl of DharmaFECT was diluted with 198 μl of optimum. Both were incubated for 5 min at room temperature (RT), mixed, and incubated for 20 min at RT. Media in each well was removed, 500 μl of the transfection media was added to each well plate with 2000 μl of antibiotic-free DMEM giving a final concentration of 25 nM of siRNA and incubated at 37°C in 5% CO2 for 48 h.

### 2.14 Determining Virus Production in XPO1 Knocked down cells

A549 cells were seeded at a density of 3.0 x 10^4^ cells per well in 6-well black-walled optical plate and incubated at 37°C in 5% CO2 for 20 h. Cells were transfected with 25 nM of siRNA targeting XPO1 or a non-targeting siRNA control and incubated at 37°C in 5% CO2 for 48 h. Cells were then infected with rLCMV/GFP-P2A-NP at an MOI of 0.03. After a 90-minute adsorption period, the inoculum was removed, and cells were treated with VC (0.5% DMSO), VE (1 µM and 5 µM), or RBV (100 µM). CCS were collected at 48 hpi, cell were washed, fixed with 4% PFA and stained with DAPI. Production of infectious particles was determined by measuring virus titers in CCS by FFA using Vero E6 cells (4 replicates) and results expressed as mean ± SD. Data were plotted using GraphPad Prism (GraphPad Software, San Diego, CA, USA). Virus infectivity was determined based on GFP expression levels quantified by fluorescence using a Biotek Cytation 5 plate reader (Agilent Technologies, Santa Clara, CA, USA). Mean values were normalized to an infected and vehicle (DMSO)-treated control, also set to 100% and plotted using GraphPad Prism v10 (GraphPad Software, San Diego, CA, USA).

### 2.15 Determining Synergistic activities

A549 cells were seeded onto a 96-well optical plate at a density of 3.0 × 10^4^ cells per well. After 20 hours, the cells were infected with rLCMV/GFP-P2A-NP MOI of 0.03. At 90 min post-infection, a concentration matrix of compounds, prepared in three-fold serial dilutions at 2x, was added in various combinations 1:1 bringing the total concentration to 1x according to a predetermined synergy matrix (8-by-8 matrix for all possible 64 combinations of two drugs combination). After 72 h post-treatment, GFP expression levels were determined. After measuring GFP, cells were subjected to DAPI staining. GFP and DAPI were quantified by fluorescence using a Biotek Cytation 5 plate reader (Agilent Technologies, Santa Clara, CA, USA). Mean values were normalized to an infected and vehicle (DMSO)-treated control, also set to 100%. Normalized values derived from matrix data were subjected to analysis utilizing the SynergyFinder+ software (https://synergyfinder.org/) [59]. Our dose-response assay generates a logistic growth fit, the drugs (included in the combination) have different mechanism of action, and is best interpreted by Zero Interaction Potency (ZIP) synergy model.[60]

## 3. Results

### 3.1. Screening of the Pandemic Response Box (PRB) for Inhibitors of LCMV Multiplication

To identify potential anti-MaAv compounds within the PRB, we implemented a HTS based on a cell-based infection assay using the rLCMV/GFP-P2A-NP, hereafter called rLCMV/GFP, that allowed us to use GFP signal as an accurate surrogate of virus multiplication[61]. To identify primary hits, we infected (MOI = 0.03) A549 cells with rLCMV/GFP and treated them with each compound at a single dose (5 µM). At 72 hpi we determined cell viability using the CellTiter 96 AQueous One Solution Cell Proliferation Assay. After absorbance values were recorded, we fixed cells using 4% PFA, and GFP expression levels quantified. Both formazan absorbance and GFP signal were determined using a Synergy H4 Hybrid MultiMode microplate reader (BioTek Instruments, Winooski, VT, USA). As controls we used treatment with the validated MaAv inhibitor ribavirin (RBV) (100 µM) and vehicle control (VC). Cell viability and GFP expression levels were normalized to those of vehicle treated (VC) controls. Primary hits (n = 22) were selected based on causing ≥ 50% reduction in infectivity (GFP signal) and ≤ 20% cytotoxicity (DAPI signal). Selected primary hits were confirmed in biological replicates, using each compound at a single concentration (5 µM) in triplicate (Fig. 1) We identified Ro-24-7429 (RO), WO 2006118607 A2 (WO), and Verdinexor (VE) as the top three hits with potent inhibitory effect on LCMV multiplication ( ≥ 80% reduction in infectivity and ≤ 20% reduction in cell viability) (Supplementary Table 1)

**Figure 1.**
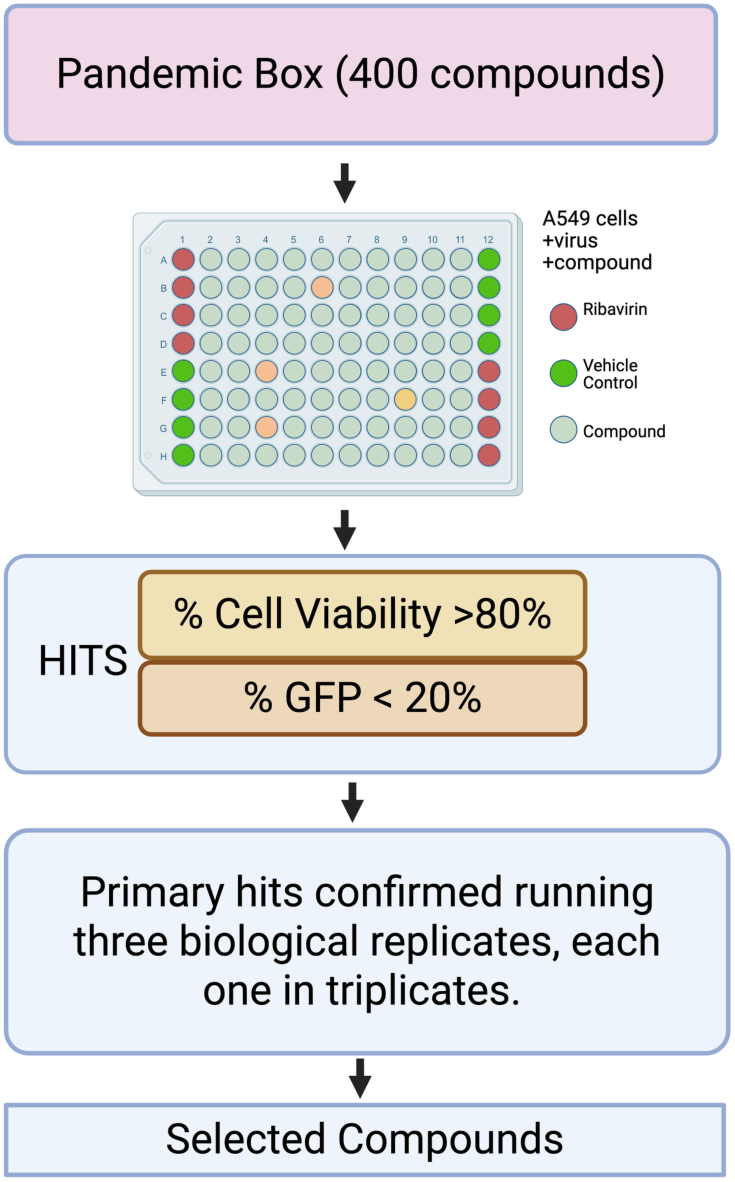
Screen Flow Chart. A549 cells were seeded (3.0 x 10^4 cells/well) in 96-well plate and 20 h later infected with rLCMV/GFP-P2A-NP at MOI of 0.03. After 90 min adsorption, the virus inoculum was removed and infected cells treated with each compound (5 µM). At 72 h pi cells were fixed with 4% PFA and stained with DAPI. GFP and DAPI staining signals were normalized to those of VC-treated and infected controls that were assigned a value of 100%. Normalized values of GFP expression and DAPI staining were used to determine infectivity and cell viability respectively. Compounds were identified as hits based on maintaining cell viability ≥ 80% and infectivity ≤ 20%. Primary hits were confirmed in biological replicates.

### 3.2. Dose-Dependent Effect of VE, RO, and WO on LCMV Multiplication in Cultured Cells

To evaluate the magnitude of the antiviral activity of the selected hits, we determined their dose-dependent effect on rLCMV/GFP multiplication in A549 cells (Fig. 2). Cells were infected with rLCMV/GFP (MOI = 0.03) and treated with threefold serial dilutions of each compound. At 72 hpi we determined cell viability using the CellTiter 96 AQueous One Solution Cell Proliferation Assay. After absorbance values were recorded, we fixed cells using 4% PFA and stained them with DAPI. GFP and DAPI staining signals, as well as formazan absorbance, were measured using the Cytation 5 Cell Imaging Multi-Mode Reader. Normalized cell viability values and GFP expression levels were used to determine compounds CC_50_ and EC_50_ values using GraphPad Prism software v10. The three selected hits VE, RO, and WO exhibited potent anti-LCMV activity with EC_50_ values (µM) of 0.074, 0.890, and 0.036 respectively, with all three having SI values > 242 (Fig. 2).

**Figure 2.**
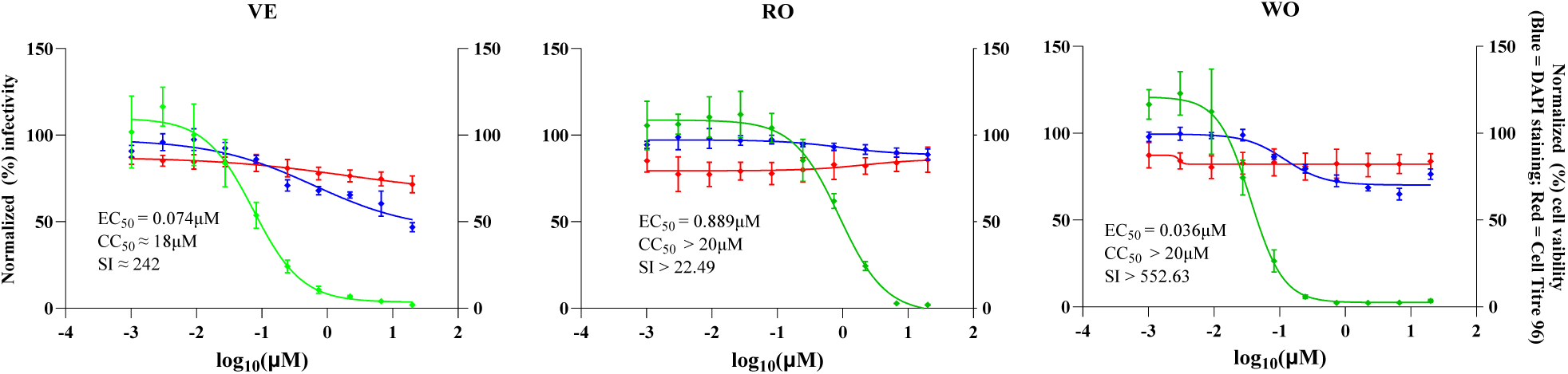
Dose-dependent effect of selected hits on LCMV multiplication. A549 cells were seeded into 96w plates (3.0 x 10^4^ cells/ well) and 20 h later infected with rLCMV/GFP-P2A-NP at MOI of 0.03. The virus inoculum was removed after 90 minutes adsorption, and infected cells treated with different concentrations (four replicates per concentration) of the indicated compounds (VE, verdinexor; RO, Ro-24-7429; WO, WO2006118607). At 72 hpi, CellTiter 96 Aqueous One solution reagent (Promega) was used to determined cell viability, the cells were washed, fixed with 4% PFA and stained with DAPI. GFP expression levels and DAPI staining signals were quantified using a Cytation 5 plate reader (BioTek, Agilent) and used to determine virus infectivity and cell viability respectively. GFP and DAPI staining values were normalized to those of VC-treated and infected cells. Mean ± SD values (n = 4 replicates) of GFP and DAPI signals were used to determine the EC_50_ and CC_50_, respectively, of each compound.

### 3.3. Efficacy of VE, RO, and WO on JUNV Multiplication in Cultured Cells

To assess the broad-spectrum anti-MaAv of VE, RO and WO, we assessed their efficacy against the LCMV genetically distantly related New World MaAv JUNV. To overcome the need of BSL4 containment required for the use of live forms of pathogenic strains of JUNV, we used rJUN GFP-P2A-NP, hereafter called rJUNV/GFP, based on the live-attenuated vaccine strain Candid#1 of JUNV, which can be safely used in BSL2. We infected (MOI = 0.5) Vero E6 (Fig.3A) and A549 (Fig.3B) cells with rJUNV/GFP and treated them with threefold serial dilution of each compound. At 72 hpi we determined cell viability using the CellTiter 96 AQueous One Solution Cell Proliferation Assay. After absorbance values were recorded, we fixed cells using 4% PFA and stained them with DAPI. GFP and DAPI staining signals, as well as formazan absorbance, were measured using the Cytation 5 Cell Imaging Multi-Mode Reader. Normalized cell viability values and GFP expression levels were used to determine compounds CC50 and EC50 values using GraphPad Prism software v10. VE and WO exhibited potent antiviral activity against JUNV in both Vero E6 (Fig. 3A) and A549 (Fig. 3B) cells, whereas RO was less efficient inhibiting multiplication of JUNV in both Vero E6 and A549 cells (Fig. 3).

**Figure 3.**
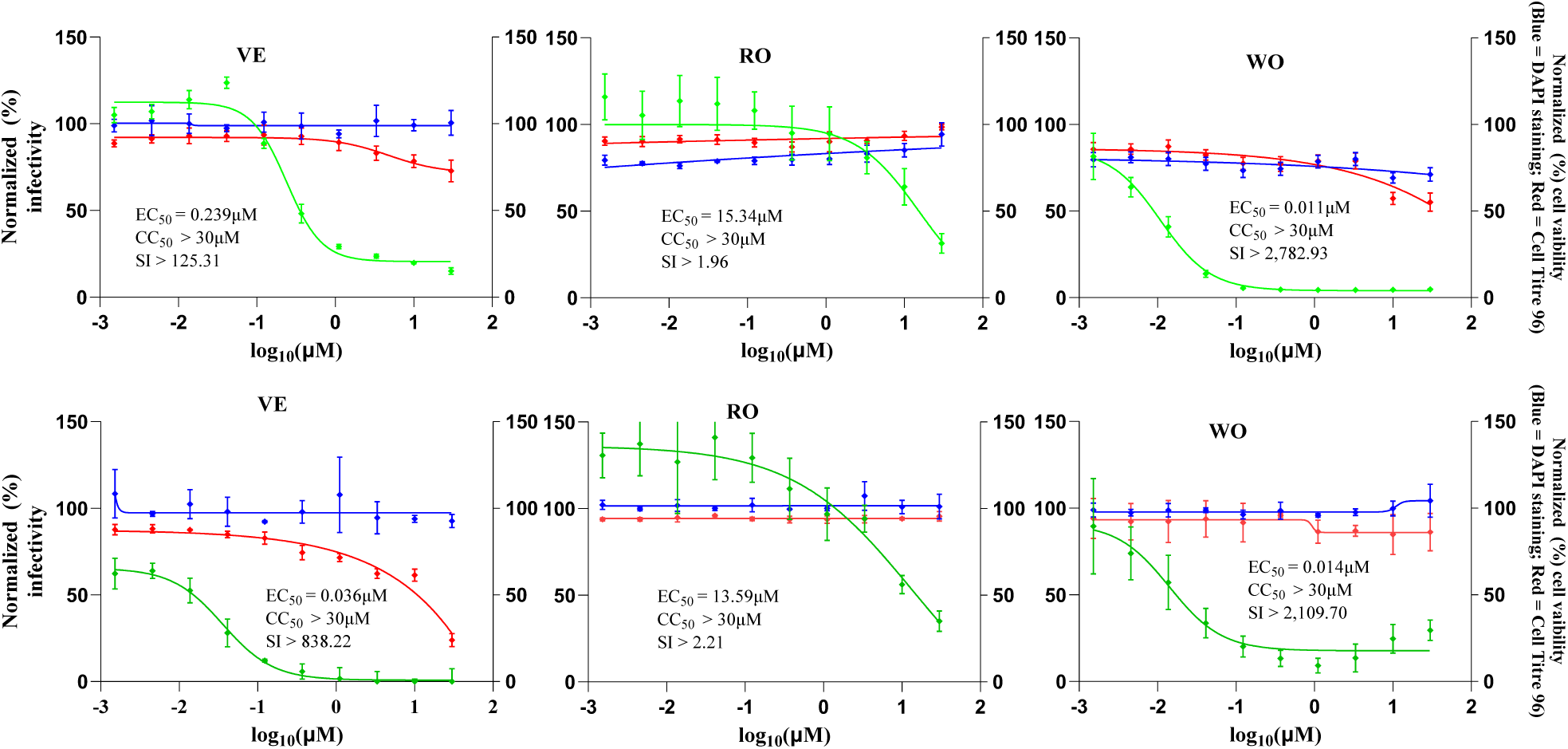
Effect of selected compounds on JUNV Multiplication in Vero E6 and A549 cells. Vero E6 (**A**) and A549 (**B**) cells were seeded into 96-well plates (3.0 x 10^4^ cells/well) and 20 h later infected with rJUNV/GFP-P2A-NP at MOI of 0.5. The virus inoculum was removed after 90-minute adsorption, and infected cells treated with different concentrations (four replicates per concentration) of the indicated compounds. At 72 hpi, CellTiter 96 Aqueous One solution reagent (Promega) was used to evaluate cell viability, the cells were washed, fixed with 4% PFA and stained with DAPI. GFP expression levels and DAPI staining were quantified using a Cytation 5 plate reader (BioTek, Agilent) and used to determine virus infectivity and cell viability respectively. GFP expression and DAPI staining values were normalized to those of VC-treated and infected cells. Mean ± SD values (n = 4 replicates) were used to determine the EC_50_ and CC_50_ respectively, of the compounds.

### 3.4. Effects of VE, RO, and WO on LCMV Multi-Step Growth Kinetics

To examine the effects of VE, RO, and WO on production of infectious viral progeny over time and virus cell propagation, we determined their effects on LCMV multi-step growth kinetics in A549 cells. We infected (MOI = 0.03) A549 cells with rLCMV/GFP and treated them with the indicated concentrations of each compound, VC, or RBV (100 µM). At the indicated hpi, CCSs were collected and titers of infectious virus determined by FFA (Fig. 4A). Cells were stained with Hoechst in FluoroBrite DMEM, and epifluorescence images of live cells were taken from each sample (Fig. 4B) using a Keyence BZ-X710 microscope. After collection of the images, total cellular RNA was isolated from each sample and levels of viral RNA synthesis determined by northern blotting (NB) (Fig. 4D) and RT-qPCR (Fig 4C). Treatment with VE at 1 µM or 5 µM caused about 2-logs reduction in virus peak titers (Fig. 4A). RO exhibited reduced efficacy at 1 µM compared to 5 µM (Fig. 4A). Treatment with WO at either 5 or 1 µM resulted in the absence of detectable levels of infectious virus in CCS at all hpi tested (Fig. 4A). The observed reduction in production of LCMV infectious progeny in CCS of infected cells treated with each of the compounds correlated with restricted viral cell propagation (Fig. 4B) and viral RNA synthesis, as quantified by RT-qPCR (Fig. 4C), or NB (Fig. 4D). VE, RO and WO inhibited to similar degree both virus replication and transcription as determined by NB as determined by NB results (Fig. 4D).

**Figure 4.**
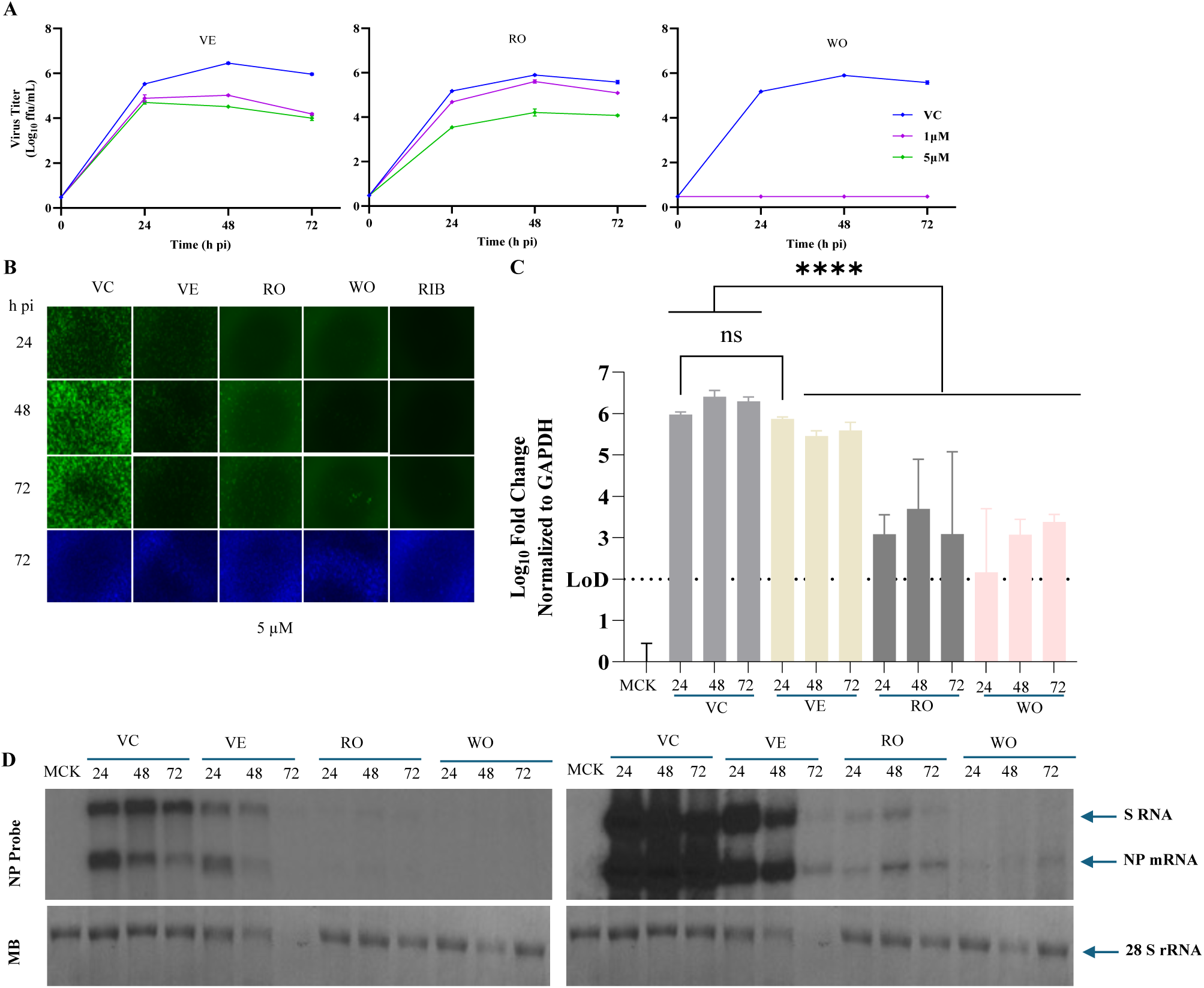
Effect of selected compounds on LCMV multi-step growth kinetics in A549 cells**. A.** Dose-dependent effect of compounds on production of infectious virus. A549 cells were seeded (2.0 x 10^5^ cells/well) in 12-well plate, and 20 h later infected with rLCMV/GFP-P2A-NP (MOI = 0.03). The virus inoculum was removed after 90-minute adsorption, and infected cells treated with VC or compounds (1 µM or 5 µM) or RBV (100 µM). At the indicated time points, cell culture supernatants (CCS) were collected, and titers of infectious virus were determined by a focus-forming assay (FFA) using Vero E6 cells. **B.** Effect of Compounds on LCMV cell propagation. At the indicated hpi, CCSs were collected, and cells were washed with PBS, stained with Hoechst dye (10µg/mL) in FluoroBrite DMEM, and images (4x magnification) were taken using a Keyence BZ-X710 microscope. **C.** Effect of compounds on viral RNA synthesis. RNA (0.5 µg) from each sample was reversed transcribed (RT), and the corresponding cDNAs used for qPCR analysis using specific primers to quantify the NP fold increase over the housekeeping gene GAPDH. A one-way analysis of variance with Šidák correction for multiple comparisons was employed for statistical analysis. Statistical significance was denoted as **** *p* < 0.0001. **D.** Effect of Compounds on virus RNA replication and transcription. After taking images of samples from B, total cellular RNA was isolated using TRI-reagent and analyzed by Northern blot using a dsDNA ^32^P probe that targets LCMV NP. Methylene blue (MB) staining was used to verify consistent transfer efficiencies across all RNA samples. Left panel shows 24 h exposure of X-ray film at 25°C, whereas the right panel shows exposure at -80°C for 24 h.

### 3.5. Effects of VE, RO, and WO on Different Steps of LCMV Life Cycle

To gain insights about the mechanisms whereby VE, RO, and WO exerted their anti-MaAv activity, we investigated their effects on various stages of the LCMV life cycle. We first assessed whether the compounds affected virus cell entry. For this, we performed a time-of-addition experiment using the single-cycle infectious rLCMVΔGPC/ZsG to prevent the confounding effects associated with multiple rounds of infection cycles in the absence of NH4Cl treatment. We treated A549 cells (3.0 × 10^4^ cells/ well, 96-well plate) with each compound (1 µM or 5 µM), the LCMV cell entry inhibitor F3406 (5 µM), or the LCMV multiplication inhibitor RBV (100 µM) starting 2 hours prior to (−2) or 2 hours post (+2) infection with rARMΔGPC/ZsG-P2A-NP at a MOI of 1. At 48 hpi we determined ZsG expression levels and normalized them to those of VC-treated cells, which were assigned a value of 100%. The results from the time of addition experiment showed that VE, RO, and WO exerted a similar inhibitory effect on ZsG expression when administered 2 h prior to or 2 h following infection (Fig. 5A), indicating that all three targeted a post cell entry step of the LCMV life cycle.

**Figure 5.**
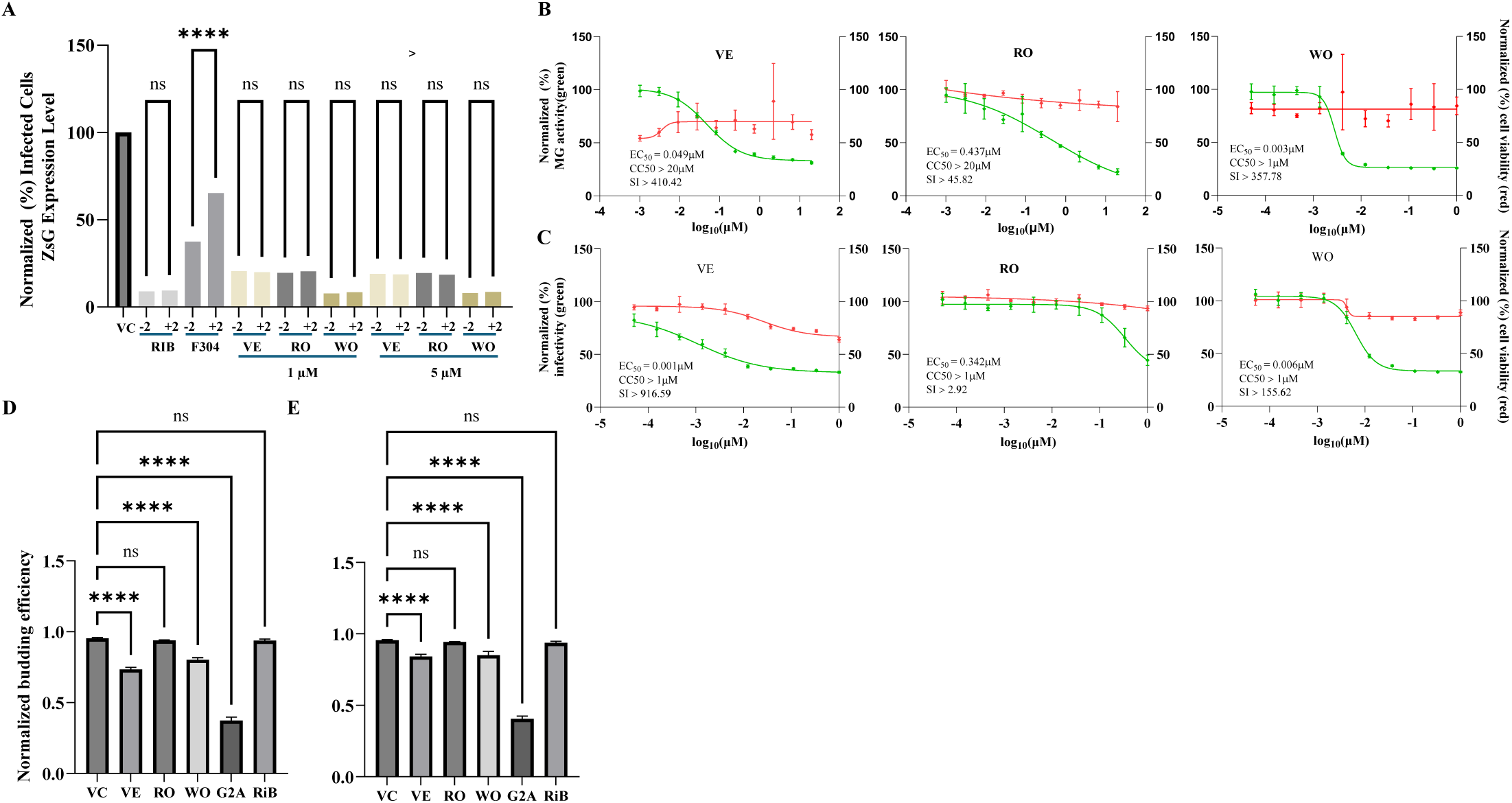
Effect of selected compounds on different steps of the LCMV life cycle. **A.** Time of addition Assay. A549 cells were seeded (3.0 × 10^4^ cells/well) in a 96-well plate and 20 h later treated with the indicated compounds at 1 µM or 5 µM, vehicle control (VC), or the LCMV cell entry inhibitor F3406 (5 µM) at either 2 h prior to (−2) or 2 h post (+2) infection with the single-cycle infectious rARMΔGPC/ZsG-P2A-NP at MOI of 1.0. At 48 hpi ZsG expression levels were determined using a Cytation 5 plate reader (BioTek, Agilent). ZsG values were normalized to those of VC-treated and infected samples, which were set to 100%. ZsG expression values from the −2 and +2 samples of each treatment were analyzed using one-way ANOVA via GraphPad Prism. **** p < 0.0001; “ns” indicates not statistically significant Effect on LASV vRNP activity using cell-based MG assay **B, C.** Cell-based MG assay. HEK293T cells were seeded (0.75 × 10^6^ cells/well) onto poly-l-lysine coated 6-well plate and 20 h later transfected with plasmids expressing the component of the LCMV (**B**) or LASV (**C**) MG system. After transfection, a single-cell suspension from each sample was prepared using Accutase, and cells seeded onto poly-l-lysine coated 96-well plate (3.0 x 10^4^ cells/well) with media containing different concentrations (four replicates per concentration) of the indicated compound. At 72 h post-transfection, cells were washed, fixed with 4% PFA and stained with DAPI. MG-directed ZsG expression levels and DAPI staining were quantified using Cytation 5 plate reader and used to determine MG activity and cell viability respectively. Mean values ± SD (n = 4 replicates) of each compound treatment normalized to VC-treated and transfected cells were used to determine the EC_50_ and CC_50_ values of the compounds. **D, E.** Budding assay. HEK293T cells were seeded (1.75 × 10^5^ cells/well) onto poly-l-lysine-coated wells in a 12-well plate, and 20 h later transfected with either pC.LASV-Z-GLuc, pC.LASV-Z-G2A-GLuc (mutant control), or pCAGGS-Empty (pC-E). At 5 h post-transfection, cells were washed three times and treated with the indicated compounds at 5 µM (**D**) and 1 µM (**E**). At 48 h post-transfection, CCSs were collected, and whole-cell lysates (WCL) prepared. GLuc activity was measured in the CCS (Z_VLP_) and WCL (Z_WCL_). GLuc activity in CCS and WCL samples were determined using the Steady-Glo Luciferase Pierce: Gaussia Luciferase Glow assay kit, and a Cytation 5 plate reader. Budding efficiency values, defined as Z_VLP_/(Z_VLP_ + Z_WCL_), were normalized (5) to those of CV-treated cells transfected with Z-WT, and analyzed (one-way ANOVA) using GraphPad Prism software (v.10). Statistical significance was denoted as **** p < 0.0001; “n” indicates a lack of statistical significance.

We next examined the effect of the compounds on the activity of the vRNP using both LCMV and LASV cell-based MG systems that accurately recapitulate all the steps of viral RNA synthesis, and where the MG-directed reporter gene expression serves as an accurate surrogate of vRNP activity. We transfected HEK293T cells with the components of the LCMV (Fig. 5B) or LASV (Fig. 5C) MG system. At 2.5 h post-transfection, single cell suspensions were prepared and seeded onto poly-l-lysine coated 96-well plate (3.0 x 10^4^ cells/well) with media containing different concentrations (four replicates per concentration) of each compound. At 72 h post-transfection, cells were fixed with 4% PFA and stained with DAPI. ZsG expression levels and DAPI staining were quantified using Cytation 5 plate reader and used to determine levels of MG activity and cell viability (total nuclei), respectively. VE and WO potently inhibited the activity of both LCMV and LASV MGs, whereas RO was significantly less efficient inhibiting either LCMV or LASV MG activities (Fig. 5B, C).

To assess the effect of VE, RO, and on the budding process directed by the matrix Z protein [62], we used a previously established cell-based budding assay wherein the activity of the Gaussia luciferase (GLuc) reporter gene serves as an indicator of Z budding activity[56]. We transfected HEK293T cells with a plasmid encoding LCMV Z-GLuc and subsequently treated them with VE, RO, WO (5 µM, and 1 µM), or VC. As controls we used the budding deficient mutant G2A of the Z [63], and treatment with RBV (100 µM). At 48 h post-transfection we measured levels of GLuc activity associated with virus-like particles (VLPs) present in CCSs, and intracellular Z protein in whole cell lysates (WCLs). Z budding efficiency was determined by the ratio of VLP-associated GLuc levels (CCSGLuc) and total GLuc levels (CCSGLuc + WCLGLuc) × 100. VE and WO, but not RO, had a significant inhibitory effect on LCMV Z budding activity at 5 µM (Fig. 5D) and 1 µM (Fig. 5E).

### 3.6. Effect of XPO1 KD on Production of LCMV Infectious progeny

VE is a selective inhibitor of the nuclear export protein XPO1 and has been shown to exhibit antiviral activity against IAV [49,64] whose replication and gene transcription take place in the nucleus, and RSV [48,65] whose M protein appears to have a nuclear function required for RSV replication. In contrast, there is no evidence that MaAv life cycle involves a nuclear phase [66], raising the intriguing question of why VE targeting of XPO1 results in MaAv inhibition. To further investigate the role of XPO1 on LCMV infection, we assessed the effect of siRNA mediated knock down (KD) of XPO1 on LCMV multiplication. For this, we transfected A549 cells with the ON-TARGETplus Human XPO1 (7514) siRNA – SMARTpool (Dharmacon) and a non-targeting siRNA as control. At 48 h post-transfection, we infected cells with rLCMV/GFP (MOI of 0.03). Non-transfected cells were treated with VE (1 µM or 5 µM), or VC. At 48 hpi CCSs were collected and titers of infectious virus determined by FFA (Fig. 6A). Cells were fixed with 4% PFA and stained with DAPI, and representative images of each sample collected using a Keyence BZ-X710 microscope (Fig. 6B). Compared to the non-targeting siRNA transfected cells, XPO1 KD in A549 cells, significantly reduced production of infectious virus (Fig. 6A), which correlated with a reduction in virus cell propagation (Fig. 6B). To confirm the KD of XPO1 in A549 cells by VE and TARGETplus Human XPO1 siRNA, we treated A549 cells with VE (5 µM), mock-treated or transfected with siRNA targeting XPO1 and non-targeting siRNA control. At 48 h post-treatment or transfection, we prepared cell lysates and levels of XPO1 protein determined by western blot (Fig. 6C). To further validate that the anti-LCMV activity of VE was via targeting XPO1, we examined the effect of the VE analog Selinexor (SE) on LCMV multi-step growth kinetics in A549 cells. SE exhibited a potent inhibitory effect on LCMV multiplication (Fig. 6D).

**Figure 6.**
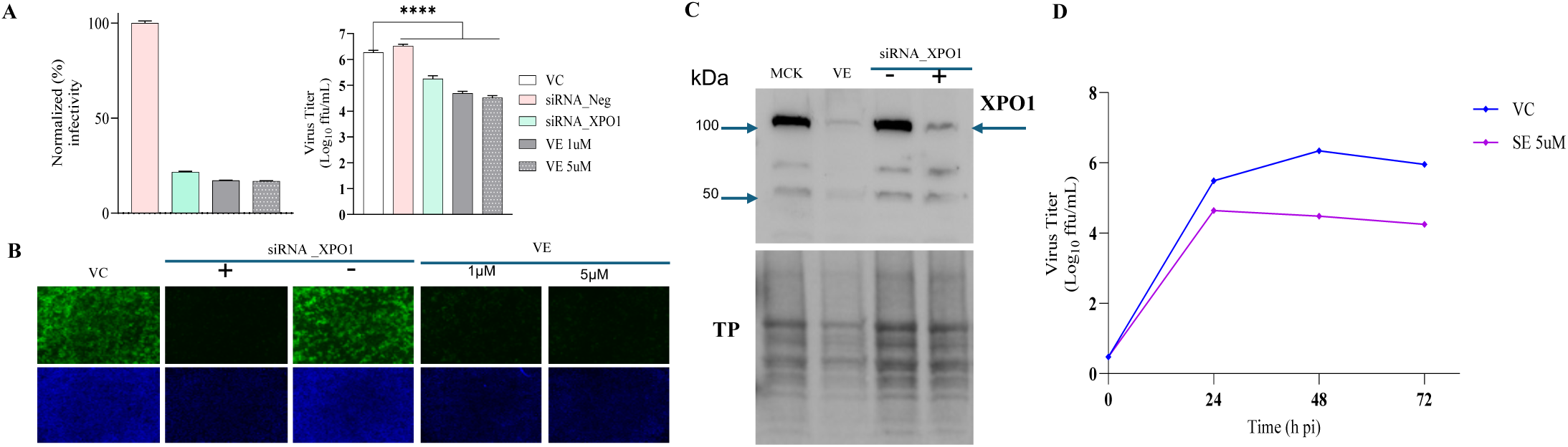
Effect of XPO1 KD on LCMV infection**. A.** Effect on production of LCMV infectious progeny. A549 cells were seeded (2.0 x 10^5^ cells/well) in 12-well plate and 20 h later transfected with siRNA targeting XPO1 or with a non-targeting siRNA control using DharmaFECT-1 reagent (Dharmacon) or left untransfected. At 48 h post-transfection, both transfected and non-transfected cells were washed and infected with rLCMV/GFP-P2A-NP (MOI of 0.03). After 90 min adsorption, the virus inoculum was removed, and cells were washed. Non-transfected cells were treated with VE (1 µM or 5 µM), or VC. At 48 hpi, CCSs were collected, and cells fixed with 4% PFA and stained with DAPI. Titers of infectious virus were determined by FFA using Vero E6 cells. GFP expression levels were quantified using a fluorescent plate reader and used to determine virus infectivity. **B.** Effect of XPO1 KD on LCMV cell propagation. Epifluorescence images (4x magnification) from A were taken using a Keyence BZ-X710 microscope. **C.** Confirmation of XPO KD. A549 cells were seeded 2.0 x 10^5^ cells/well) in 12-well plate and 20 h later treated with VE (5 µM), mock-treated or transfected with siRNA targeting XPO1 or a non-targeting siRNA control using DharmaFECT-1 reagent (Dharmacon). At 48 h post-treatment, cell lysates were prepared and XPO1 protein expression examined by western blotting. **D.** Dose-dependent effect of SE on production of infectious virus. A549 cells were seeded (2.0 x 10^5^ cells/well) in 12-well plate, and 20 h later infected with rLCMV/GFP-P2A-NP (MOI = 0.03). The virus inoculum was removed after 90-minute adsorption, and infected cells treated SE (1 µM or 5 µM) or VC. At the indicated time points, cell culture supernatants (CCS) were collected, and titers of infectious virus were determined by a focus-forming assay (FFA) using Vero E6 cells.

### 3.7. Assessment of the Contribution of the Interferon (IFN) Response to VE Anti-MaAv Activity

Treatment with VE and the related second generation of SINE Eltanexor has been shown to increase expression of type I interferon (T1IFN) and other innate immune-related genes in Kaposi’s sarcoma-associated herpesvirus (KSHV) and Human Cytomegalovirus (HCMV) infections [67,68], whereas treatment with VE and SE has been shown to decrease levels of type II IFN and other proinflammatory cytokines in IAV and SARS-CoV-2 infections [64,69]. To assess the contribution of the T1IFN response to the anti-MaAv we evaluated the dose response effect of VE on LCMV multiplication in A549 cells deficient in mitochondrial antiviral-signaling protein (MAVS) required for induction of IFNb in LCMV-infected cells[70]. VE exerted a strong dose-dependent inhibitory effect on LCMV multiplication in MAVS-KO A549 cells (Fig.7A), which correlated with reduced production of infectious progeny (Fig. 7B) and restricted virus propagation in the infected cell monolayer (Fig. 7C). These findings indicated that the T1IFN response did not significantly contributed to the anti-LCMV activity of VE. Unexpectedly, we found that under our experimental conditions, treatment with VE resulted in reduced expression levels of the interferon stimulated genes ISG15 and MX1 (Fig. 7D).

**Figure 7.**
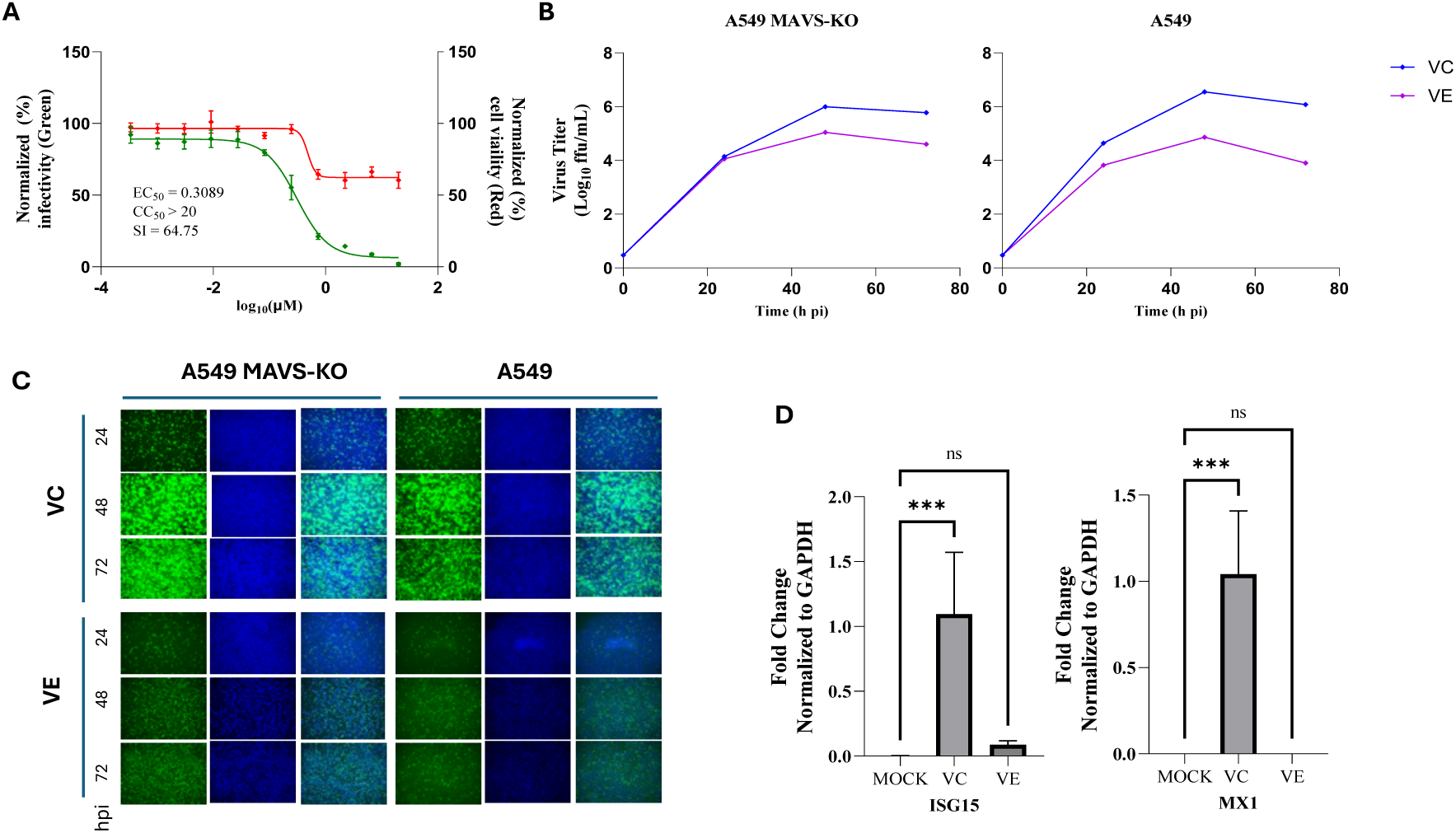
Type 1 interferon does not contribute to VE anti-MaAv activity. **A**. Dose-dependent inhibitory effect of VE on LCMV multiplication in A549 MAVS-KO cells. Cells were seeded into 96-well plates (3.0 x 10^4^ cells/ well) and 20 h later infected with rLCMV/GFP-P2A-NP at MOI of 0.03. The virus inoculum was removed after 90 minutes adsorption, and infected cells treated with different concentrations (four replicates per concentration) of VE. At 72 hpi cells were fixed with 4% PFA and stained with DAPI. GFP expression levels and DAPI staining were quantified using a Cytation 5 plate reader (BioTek, Agilent). GFP expression and DAPI staining values were normalized to those of VC-treated and infected cells. Mean ± SD values (n = 4 replicates) were used to determine the EC_50_ and CC_50_ of the compounds. **B.** Effect of VE on LCMV multi-step growth kinetics in A549 MAVS-KO cells. A549 WT and MAVS-KO cells were seeded (1.5 x 10^5^ cells/well) in 24-well plates, and 20 h later infected with rLCMV/GFP-P2A-NP (MOI = 0.03). The virus inoculum was removed after 90-minute adsorption, and infected cells treated VC, VE (5 µM), or RBV (100 µM). At the indicated time points, CCSs were collected and virus titers determined by FFA using Vero E6 cells. **C.** Effect of VE on LCMV cell propagation. At the indicated hpi, cells samples from **B** were were fixed with 4% PFA and stained with DAPI. Representative images (4x) were taken using a Keyence BZ-X710 microscope. **D.** Effect of VE the expression of ISGs ISG15 and MX1 in LCMV-infected A549 cells. A549 cells were seeded (2.0 x 10^5^ cells/well) in 12-well plate, and 20 h later infected with rLCMV/GFP-P2A-NP (MOI = 0.03). The virus inoculum was removed after 90-minute adsorption, and infected cells treated VE (5 µM), or VC. At 48 hpi, CCS was removed, cells were washed with PBS and total cellular RNA was isolated from each sample using TRI-reagent. RNA (0.5 µg) from each sample was reversed transcribed (RT), and the corresponding cDNAs used for qPCR analysis to quantify expression levels of the ISG15 and MX1 genes with normalization to the housekeeping gene GAPDH. A one-way analysis of variance with Dunnett correction for multiple comparisons was employed for statistical analysis. Statistical significance was denoted as *** *p* < 0.001.

### 3.8. Assessment of the Synergistic MaAv Antiviral Activity of VE in Combination with WO and RO

To investigate the synergistic antiviral activity of VE with WO and RO against MaAv, we designed an 8x8 matrix to test 8 serial dilutions of two compounds in their 64 possible combinations for their ability of inhibiting multiplication of LCMV. VE showed strong synergistic antiviral activity in combination with WO (Fig. 8A) with highest ZIP synergy score of 20 observed at concentrations of 0.11μM (VE) and 0.0003 μM (WO). Monotherapy treatment with VE and WO at these concentrations resulted in moderate antiviral effect of VE (22.64% inhibition) and non-detectable activity for WO. However, when combined, VE+WO exhibited 50.05% inhibition. Cell viability with individual compound treatment at these concentrations was 69.57% (VE) and 81.24% (WO), whereas combination therapy resulted in 72.72% cell viability. The combination treatment of VE and WO also showed five other combinations with ZIP score values > 10. The combination therapy of VE and RO only exhibited very modest synergistic antiviral activity with highest Zip synergy score of 7.6 (Fig. 8B) observed at concentrations of 0.333μM (VE) and 1.333μM (RO). The individual treatments of VE and RO at these concentrations displayed moderate antiviral effects with percent inhibition value of 77.62% and 40.14% respectively. However, when combined, 96.24% inhibition was exhibited. Cell viability with individual treatment at same concentration was 53.41% and 100%, respectively, while cell viability when in combination was 54.3%. The combination treatment of VE and RO also showed four other combinations with ZIP score values > 5.

**Figure 8.**
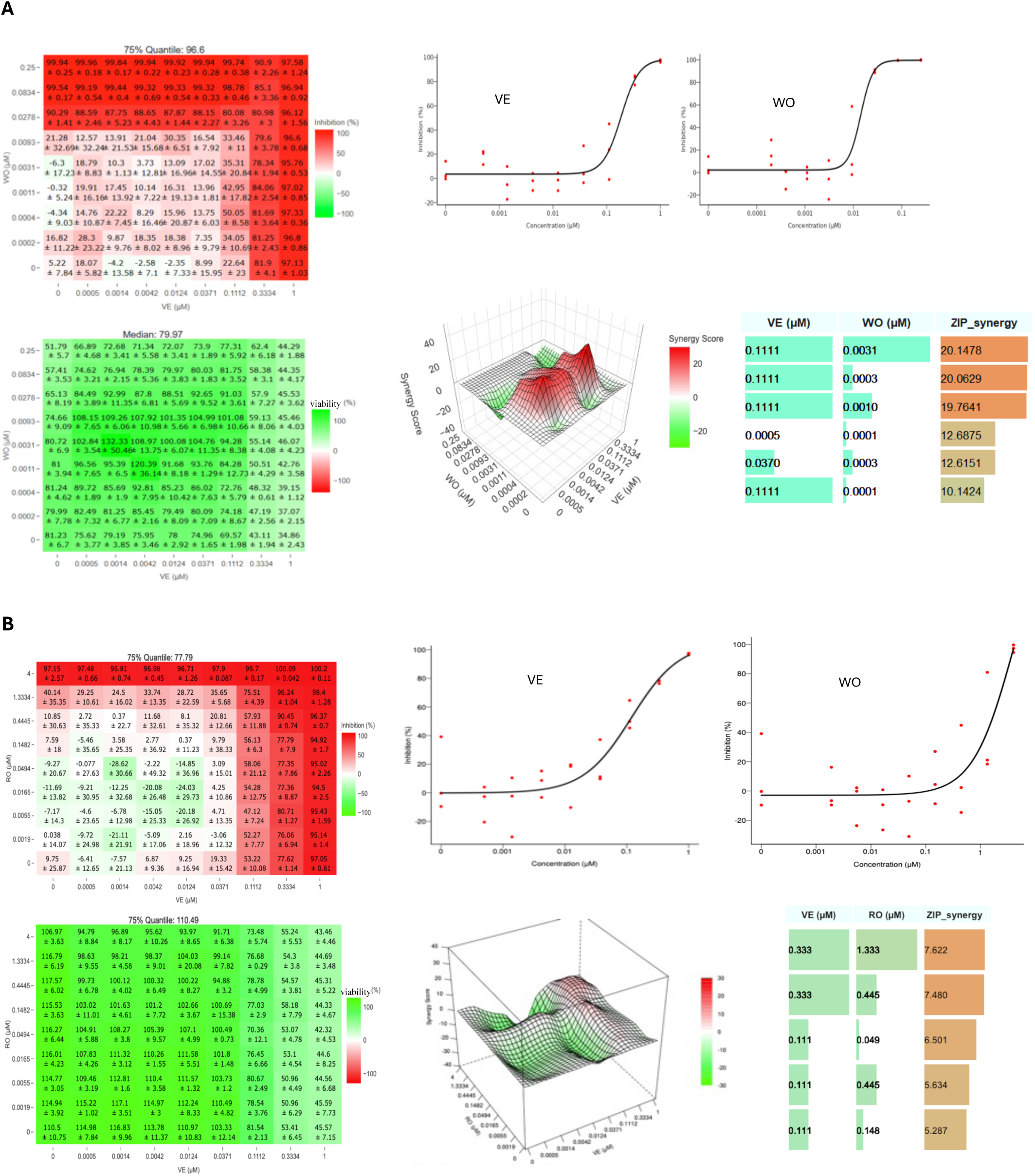
Synergistic antiviral activity of VE in combination with WO (**A**) and RO (**B**). A549 cells were seeded at 3 *×* 10^4^cells/well into 96-well plates, and 20 h later infected rLCMV/GFP-P2A-NP at MOI of 0.03. After 90 min adsorption, the virus inoculum was removed and medium containing the indicated drug combinations and concentrations added to the cells. At 72 hpi, the cells were washed, fixed with 4% PFA and stained with DAPI. GFP expression levels and DAPI staining were quantified using a Cytation 5 plate reader (BioTek, Agilent). GFP and DAPI values were normalized to those of VC-treated and infected cells. The data was uploaded into the software using the table template. The % inhibition option for GFP signal and % viability option for DAPI were used for the response specification step of the software. The synergy matrices’ heat maps, dose–response curves, and three-dimensional synergy plots and scores were generated using the SynergyFinder+ software. Bar plots of the synergistic dose–response were obtained using RStudio software (version2025.09.1+401).

## 4. Discussion

The lack of US FDA-approved vaccines or antivirals to prevent and treat diseases caused by human pathogenic MaAv represents an important unmet medical need. Current treatment of infections by LASV and other human pathogenic MaAv is limited to an off-label use of ribavirin whose efficacy remains controversial[17,18,71]. Significant efforts are being directed towards the discovery and development of antiviral drugs against LASV and other hemorrhagic fever causing MaAv, which has resulted in the identification of several promising candidates [35]. However, expanding the landscape of MaAv candidate antivirals will facilitate overcoming some current challenges including limited potency and the development of resistance. Drug-repurposing strategies can accelerate the progression of a candidate antiviral drug into clinical trials, thereby reducing the labor-and resource-intensive efforts associated with the preclinical optimization of newly discovered hits in traditional drug-discovery approaches [34,72].The open-source PRB is a collection of 400 compounds assembled based on publicly available information regarding chemotypes currently in the discovery and early development stages, which have demonstrated beneficial activities [39].

Compounds from the PRB have been identified as potential antivirals against different viruses including enterovirus A71[73], KSHV[74], SARS-CoV-2[75,76], Ebola virus [77], Chikungunya virus[78], and Zika virus[79]. Here we have presented our results from the screening of the PRB for compounds with anti-MaAv activity. We used a cell-based infection assay with features amenable for HTS [57,61] to screen the PBR for inhibitors of LCMV multiplication. We identified 22 primary hits. Validation and reassessment assays identified RO, WO and VE as the top candidate inhibitors of LCMV multiplication in cultured cells. RO (Ro-24-7429) was originally developed as an inhibitor of the HIV-1 Tat protein [40,41] but with no detectable antiviral effect in HIV-infected patients [42] [80]. Subsequently, RO was found to inhibit runt-related transcription factor 1(RUNX1) protein[43], and to ameliorate lung fibrosis and inflammation in a mouse model of bleomycin-induced pulmonary fibrosis[43]. Additionally, RO was shown to reduce the expression of angiotensin-converting enzyme 2 (ACE-2) and furin, which are host proteins essential for SARS-CoV-2 infection, supporting the repurposing of RO for the treatment of SARS-CoV-2 infection[43]. We found that RO exhibited strong antiviral activity against LCMV (EC50 = 0.89 µM and SI >22). RO did not affect viral cell entry or Z-mediated budding processes, but rather inhibited RNA synthesis mediated by vRNP (Fig. 4D, and Fig. 5). However, unexpectedly, RO showed significantly lower activity against JUNV (EC50 =15.34 µM and SI = 1.96), questioning its use as broad-spectrum MaAv antiviral. WO, a tetrahydrocarbazole amide, has been shown to exhibit broad-spectrum antiviral activity[81], which has been linked to its inhibitory effect on dihydroorotate dehydrogenase (DHODH)[82] a key enzyme in the biosynthesis of pyrimidines [44,45]. WO showed potent antiviral activities against LCMV (EC_50_ = 0.04 µM and SI >552.63) and JUNV (EC_50_ = 0.01µM and SI >2.8) via targeting the activity of the vRNP, a result consistent with our published findings showing that DHODH inhibitors brequinar[83] and vidofludimos calcium and related analogs [84] have potent anti-MaAv activity.

VE, an oral Selective Inhibitor of Nuclear Export (SINE) compound that targets the CRM1/XPO1 exportin [46] has been shown to inhibit the replication of IAV [47] and RSV [48] in cultured cells. Notably, VE has demonstrated efficacy in animal models of IAV infection [47,49]. VE inhibits IAV multiplication by blocking XPO1-mediated nuclear export of vRNP [47], whereas VE induced nuclear accumulation of RSV M protein due to of inhibition of its XPO1-mediated nuclear export of RSV M protein is the primary mechanism by which VE and other SINE compounds inhibit RSV multiplication in cultured cells [48]. VE has also been shown to have antiviral activity against several DNA viruses via nuclear retention of viral proteins and also promoting the production of type I interferon [85,86]. MaAv life cycle is restricted to the cytoplasm of infected cells [66]. Hence, it was unexpected to find that VE exhibited a potent antiviral effect against MaAv, with EC_50_ values of 0.07 µM and 0.24 µM against LCMV and JUNV, respectively, and corresponding SI values of 242 and 124.31. We found that siRNA mediated KD of XPO1 phenocopy the effect of VE in LCVM-infected cells, characterized by XPO1 reduced expression levels that correlated with inhibition of production of LCMV infectious progeny (Fig. 6), supporting that VE exerted its anti-MaAv activity via targeting XPO1. Increased expression of T1IFN has been implicated in inhibition of KSHV and HCMV infection by treatment with SINE compounds [68,86]. However, we found that VE exhibited an anti-LCMV activity of similar magnitude in WT and MAVS-KO A549 cells, indicating that the host cell T1IFN response did not contribute significantly to the antiviral effect of VE on LCMV. Moreover, contrary to the reported VE induced expression of ISGs in KSHV and HCMV infected cells [67,68], we observed a decreased expression of ISG15 and MX1 ISGs upon VE treatment (Fig. 7). Recent evidence suggest that XPO1 has activities beyond its role in nuclear export including epigenomic regulation[87], which could influence the host cell response to infection.

HDAs represent a promising yet underexplored approach in contemporary antiviral research [88,89]. Unlike conventional direct-acting antivirals (DAAs) that target viral proteins and functions, HDAs disrupt host factors and cellular processes that viruses hijack for their replication and pathogenesis. Since members of a virus family share host dependencies, HDAs have the potential to act as broad-spectrum antivirals. In addition, HDAs pose a higher genetic barrier to the emergence of drug-resistant viral variants, which often compromise antiviral therapy[50,90]. The current fast pace of identification and characterization of virus-host interactions has accelerated the discovery of many novel potential host druggable targets for the development of HDAs and have facilitated drug repurposing strategies for drugs with established safety profiles, thus reducing longer development timelines and regulatory barriers associated with the development of novel drugs[91]. HDA-based antiviral strategies face the challenge of risks associated with toxicity and unintended consequences of disruption of cell physiological processes. However, therapies against acute viral infections, such as HF diseases causes by MaAv, require short-term treatment, which increases the likelihood of identifying effective therapeutic regimens with acceptable safety profiles.

The success of combination therapy is illustrated by the current therapeutic approaches to treat HIV and HCV infections involving combinations of two to four antivirals targeting different steps of the virus life cycle [92–95]. This has the advantage of fostering synergistic antiviral effect [96,97], which facilitates reducing drug doses within therapeutic range and alleviate side effects associated with high drug doses using in monotherapy, as well as posing a higher genetic barrier to the emergence of drug-resistant viral variants that often jeopardize monotherapy [98,99]. We investigated the synergistic efficacy of VE with RO and WO and found that VE exhibited strong synergistic antiviral activity in combination with WO (Fig. 8A) with ZIP synergy score of 20. In contrast, the combination therapy of VE and RO only exhibited very modest synergistic antiviral activity with Zip synergy score of 7.6 (Fig. 8B). Future studies should examine whether VE exhibits synergistic antiviral activity in combination with inhibitors of LASV cell entry that have been advanced to clinical trials[100], which could facilitate the development of combination therapies of HDAs and DAAs. It should be noted that research aimed at developing VE, and related XPO1 targeting SINE compounds, as a host-directed antiviral against MaAv could be facilitated by insights gathered from cancer research, including preclinical and clinical data from the oncology pipeline for XPO1 inhibitors[101].

## Supplementary Materials

The following supporting information can be downloaded at: https://www.mdpi.com/article/doi/s1, Figure S1: title; Table S1: title; Video S1: title.

## Author Contributions

For research articles with several authors, a short paragraph specifying their individual contributions must be provided. The following statements should be used “Conceptualization, C.A.O., B.C. and J.C.T.; methodology, C.A.O., B.C., H.W. and J.C.T.; software, C.A.O. and H.W.; validation, C.A.O., B.C., H.W. and J.C.T.; formal analysis, C.A.O. and H.W.; investigation, C.A.O. and B.C.; resources, N.O. and J.C.T..; data curation, C.A.O., B.C. and C.B.O.; writing—original draft preparation, C.A.O.; writing—review and editing, C.A.O., B.C., H.W. and J.C.T.; visualization, C.A.O. and B.C.; supervision, H.W. and J.C.T.; project administration, C.A.O. and B.C.; funding acquisition, J.C.T. All authors have read and agreed to the published version of the manuscript.” Please turn to the <u>CRediT taxonomy</u> for the term explanation. Authorship must be limited to those who have contributed substantially to the work reported.

## Institutional Review Board Statement

No applicable.

## Informed Consent Statement

No applicable.

## Data Availability Statement

The raw data supporting the conclusions of this article will be made available by the authors on request.

## Acknowledgments

No applicable

## Disclaimer/Publisher’s Note

The statements, opinions and data contained in all publications are solely those of the individual author(s) and contributor(s) and not of MDPI and/or the editor(s). MDPI and/or the editor(s) disclaim responsibility for any injury to people or property resulting from any ideas, methods, instructions or products referred to in the content.

